# High Yield Monolayer Graphene Grids for Near-Atomic Resolution Cryo-Electron Microscopy

**DOI:** 10.1101/827089

**Authors:** Yimo Han, Xiao Fan, Haozhe Wang, Fang Zhao, Christopher G. Tully, Jing Kong, Nan Yao, Nieng Yan

## Abstract

Cryogenic electron microscopy (cryo-EM) has become one of the most powerful techniques to reveal the atomic structures and working mechanisms of biological macromolecules. New designs of the cryo-EM grids—aimed at preserving thin, uniform vitrified ice and improving protein adsorption—have been considered a promising approach to achieving higher resolution with the minimal amount of materials and data. Here, we describe a method for preparing graphene cryo-EM grids with 99% monolayer graphene coverage that allows for more than 70% grid squares for effective data acquisition with improved image quality and protein density. Using our graphene grids, we have achieved 2.6 Å resolution for streptavidin, with a molecular weight of 52 kDa, from 11,000 particles. Our graphene grids increase the density of examined soluble, membrane, and lipo-proteins by at least five times, affording the opportunity for structural investigation of challenging proteins which cannot be produced in large quantity. In addition, our method employs only simple tools that most structural biology laboratories can access. Moreover, our approach allows for customized grid designs targeting specific proteins, due to its broad compatibility with a variety of nanomaterials.

**Significance statement:** Single particle cryogenic electron microscopy (cryo-EM) represents the cutting-edge technology to determine three-dimensional atomic structures of bio-macromolecules. However, issues of cryo-sample preparation limit the cryo-EM to achieve higher resolution. Here, we demonstrated a high yield, monolayer graphene supporting film to improve the cryo-sample quality. Using our approach, we have achieved so far, the highest resolution structure of the smallest protein by cryo-EM with the minimal number of datasets. Our technique paves the way for universal cryo-sample preparation for near-atomic resolution cryo-EM.

## Introduction

Cryogenic electron microscopy (cryo-EM) provides an effective way to investigate the structures of biological macromolecules ^1–4^. Technology breakthrough in direct electron detection ^5,6^ and advanced algorithms^6,7,8^ have enabled cryo-EM to map the precise structural details of biological macromolecules at near-atomic resolutions ^9^, which is essential for the understanding of their functions. As the cryo-EM community expands, it is a share view of many researchers that the bottleneck for cryo-EM resides in sample preparation. Cryo-EM requires protein particles to be suspended in a thin layer of vitrified ice to avoid denaturation ^10,11^. To achieve this, continuous amorphous carbon films and holey carbon grids (Quantifoil) have been widely used. Followed by glow-discharge plasma treatment, these grids become hydrophilic and allow the formation of a thin layer of aqueous solution upon blotting by a filter paper ^12^. Among these grids, the continuous carbon film (usually 20 nm thick) inevitably introduces electron scattering that adds noise and reduced image resolution. Therefore, holey carbon grids, where the layer of solution can form in the hole area, have been considered the preferred cryo-EM grids for high resolution single particle analysis. However, due to the distinct protein properties, for which we coined a term *proteinalities*, Quantifoil grids do not work for all proteins. While some proteins prefer to attach to the carbon film and fail to enter the holes, others stay at the air-water interface with compromised folding ^13^. In addition, the non-uniformity of ice thickness makes it difficult to search across the entire grids for thin ice area, where the image contrast is optimal for high-resolution image processing. Since thin ice and high protein density are key to high-resolution reconstruction of the protein structure, a better design of the cryo-EM grid that can solve these problems will benefit the cryo-EM community.

Graphene materials (such as graphene oxide (GO)) have been used as supporting films in cryo-EM to improve the protein density in the hole areas of Quantifoil grids ^14,15,16,17^ (Schematically shown in Fig. 1a and 1d). Compared to films made by other materials, graphene has the advantages of being intrinsically thin and made of only light elements (carbon, oxygen, hydrogen, etc). These advantages make functional graphene transparent to 300 kV electrons. Among the graphene materials, GO films have been tested using 20S proteasome (700 kD) at a low concentration (tens of microgram per milliliter) to reconstruct a structure at ~2.5 Å resolution with reasonable adsorption ^18^. Despite the improvements of protein adsorption, these grids are hardly used in the community mainly for three reasons: (1) low coverage of graphene limits the effective areas to acquire cryo-EM data; (2) non-uniform surface contamination results in either protein aggregation or no adsorption; (3) intricate fabrication process or requirement of expensive instruments that most structural biology labs have no access to.

**Figure 1.**
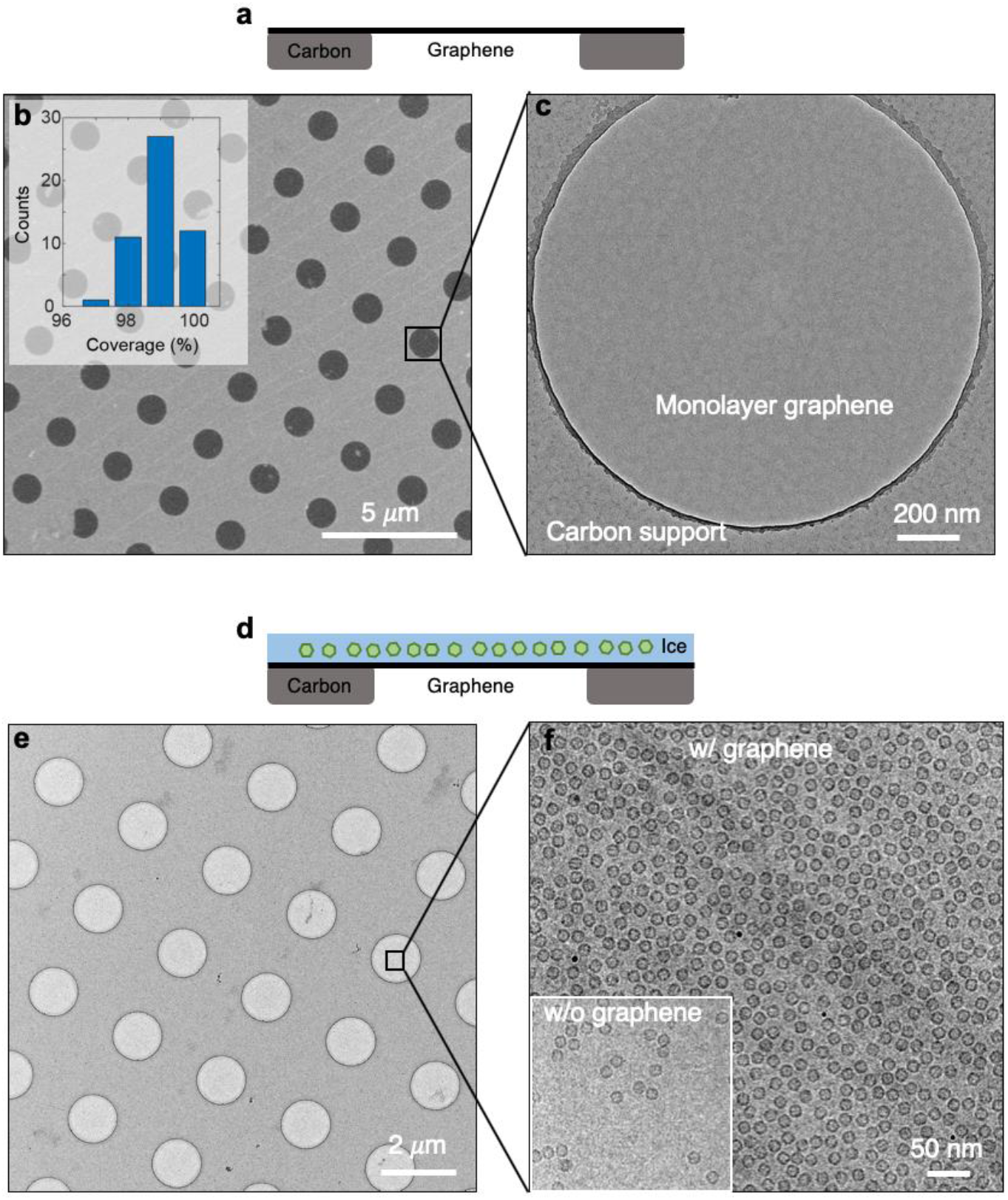
High quality graphene grids for cryo-EM. **a,** Side-view schematic of graphene grids. **b,** Large-scale SEM image of graphene film on holey carbon TEM grids. All the holes in this area are covered by suspended graphene. The inset displays the statistics of graphene yield. In average, 99% of graphene has been successfully suspended over holes. The statistics were conducted by counting the yield of suspended graphene in 50 squares. **c,** A zoomed-in TEM image of suspended graphene, showing its uniformity and cleanness. **d,** Side-view schematic of cryo-EM sample on graphene grids. **e,** Low-magnification image of a cryo-EM sample using graphene grids. The uniformity and cleanness of graphene contributes to a uniform and thin ice layer with proteins embedded. **f,** Cryo-EM micrograph of apoferritin on graphene grids, compared to the same sample on holey carbon grids (inset). The apoferritin concentration in solution is 1.2 mg/mL.

Here, we demonstrate a more convenient and less costly approach to fabricating high-quality graphene cryo-EM grids with nearly full graphene coverage (Fig. 1b) and clean graphene surfaces (Fig. 1c), which provide uniform and thin ice layer (Fig. 1e) and improve the protein density (Fig. 1f) for single particle cryo-EM analysis.

## Results

### Fabrication of graphene grids

We fabricated graphene cryo-EM grids by transferring continuous monolayer graphene from its original substrate (copper) to a Quantifoil holey carbon grid using an organic molecule assisted transfer method, as schematically described in Fig. 2a (more details in Supplementary Fig. 1). With a thin layer of Methyl methacrylate (MMA) to support graphene during the transfer process, our method results in a very high percentage coverage of hole areas by suspended graphene. Fig. 1b shows an example where all holes in the holey carbon film are covered without any broken one. Statistics from different areas indicate that the average yield of suspended monolayer graphene is ~99% (inset of Fig. 1b), higher than any previously reported functional graphene cryo-EM grids. In addition, our cleaning process is sufficient to remove most organic molecule residues and achieve clean graphene surface (Fig. 1c). The entire fabrication process takes about a few hours, and the production of up to hundreds of grids can be done en masse without special requirement of equipment or large quantity of reagent (an overview of the method in Supplementary Fig. 1 and a video demonstration in Supplementary Video).

**Figure 2.**
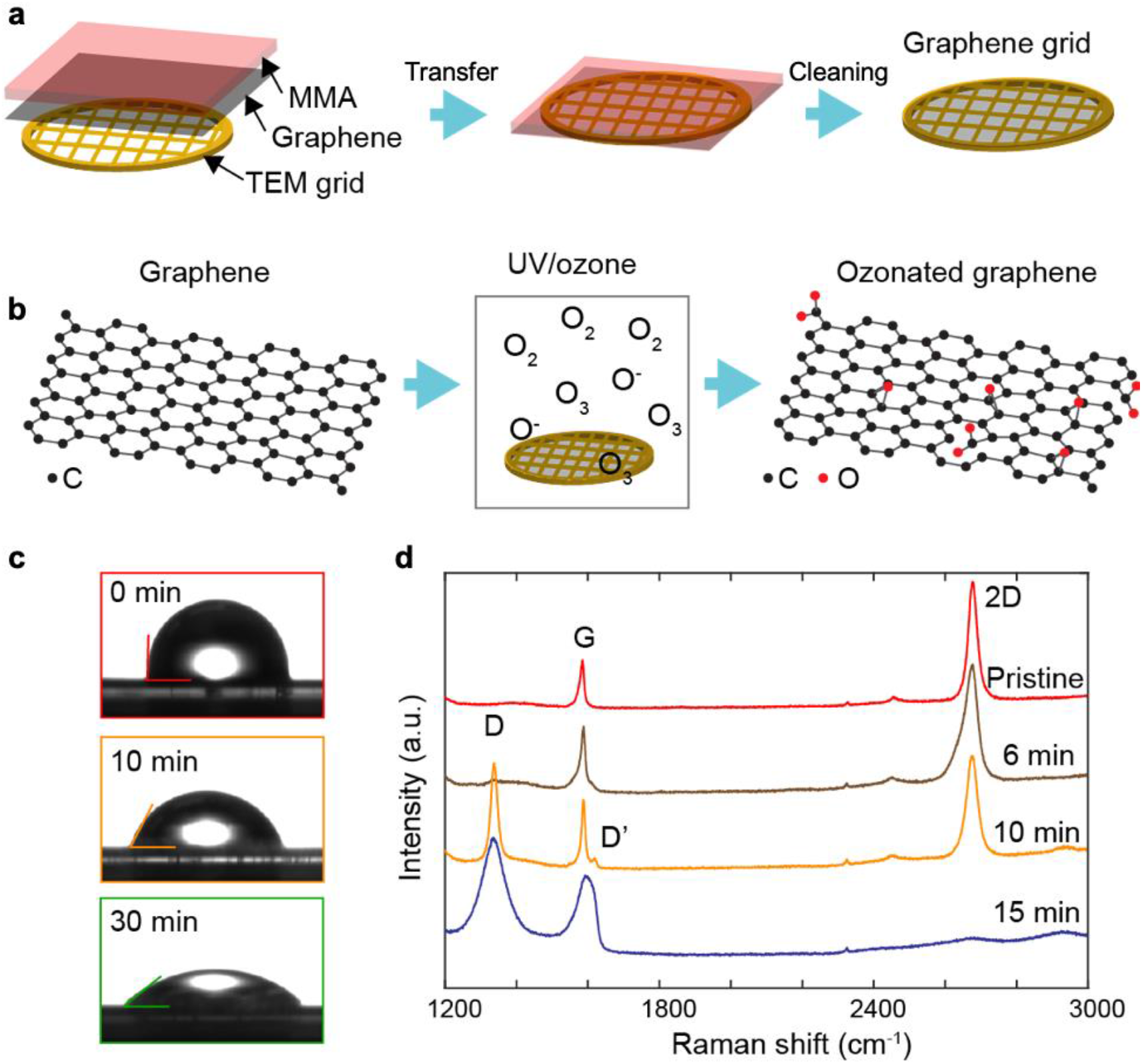
Fabrication of graphene cryo-EM grids. **a,** Schematic summary of graphene grid fabrication process. More details are provided in Supplementary Fig. 1. **b,** Schematic of surface treatment by UV irradiation. The ozonation of graphene adds oxygenated functional groups to the graphene surface, which makes graphene hydrophilic. **c,** Contact angle measurements showing that UV/ozone increase the hydrophilicity (original graphene: 89.4°; 10 min: 71.6°; 30 min: 45.5°). **d,** Raman spectroscopy shows that 10-min UV/ozone treatment can convert graphene to oxygenated graphene by creating defects (indicated by the D peak in the orange curve). 15 min UV/ozone can effectively convert graphene into GO (blue curve).

Since graphene is hydrophobic, their natural surfaces show limited adsorption of proteins (Supplementary Fig. 2). Conventional glow-discharge plasma cleaning (Ar or O_2_ plasma) that has been widely used to generate hydrophilic carbon films will damage graphene in seconds, owing to the atomic thinness of graphene. Instead, we employed UV/ozone, which uses UV irradiation to generate a small amount of ozone gas to gently oxidize sample surfaces (Fig. 2b). UV/ozone has been widely used to clean the surface of semiconductors and polymers ^19,20^. The ozonation of monolayer graphene forms oxygenated functional groups, which can effectively render the surface of graphene hydrophilic ^21–23^.

UV/ozone has the advantage of adding functional groups to graphitic surface at a slow and thus controllable rate, therefore fine-tuning the surface properties of graphene. The contact-angle measurements (Fig. 2c) demonstrate that graphene gradually becomes hydrophilic as the UV exposure time increases. Within ten minutes, UV/ozone effectively converts graphene into hydrophilic graphene, as shown in the orange box in Fig. 2c. Raman spectroscopy (Fig. 2d) indicates the hydrophilic graphene after 10 min UV irradiation is composed of oxygenated graphene (orange curve in Fig. 2d), while five more minutes UV irradiation turns graphene into complete GO (blue curve in Fig. 2d). X-ray photoelectron spectroscopy (XPS) analysis (Supplementary Fig. 3a) indicates an increment of -COO groups and C-O bonds from the oxygenated functional groups. In the XPS plots, a noticeable drop of C-O peak appears in the sample in UV irradiation after ten minutes, indicating that a cleaning process removes the surface contaminants on graphene due to air exposure. In addition, we found that under UV irradiation, graphene films can survive up to an hour, where nanopores start to emerge and expand (Supplementary Fig. 3b). As the ten-minute treatment is relatively gentle and presents clean and uniform surfaces with good hydrophilicity, we used this parameter to treat our graphene for the following cryo-EM experiments.

### A 2.2 Å resolution reconstruction of apo-ferritin with graphene grids

To demonstrate that, practically, our high-quality graphene grids are suitable for cryo-EM at near atomic resolution, we prepared standard apo-ferritin on our graphene grids for cryo-EM data collection and single particle analysis (Fig. 1f). In addition to the improved protein density, we have reached a high-resolution reconstruction of apo-ferritin at 2.2 Å (Fig. 3a and Supplementary Fig. 4a). The resolution was determined by gold standard Fourier shell correlation (FSC) (Supplementary Fig. 4b), where the information limit has already reached the Nyquist frequency of the input micrographs, indicating our graphene grids are suitable for resolution beyond 2.2 Å. In the reconstructed local density maps, we were able to clearly dock individual residues in the side chain from the published PDB model (Fig. 3b). In addition, the central holes of the benzene rings were clearly resolved in the density map (Fig. 3b, Phe_51_ and Tyr_137_), confirming a veritable high resolution we have achieved using the graphene grids. Our reconstruction of apo-ferritin at 2.2-Å resolution is the highest among those in EM databank (EMDB) using GO or other thin film supported grids. These results indicate the potential of our graphene grids in cryo-EM investigations of protein structures at a near-atomic resolution.

**Figure 3.**
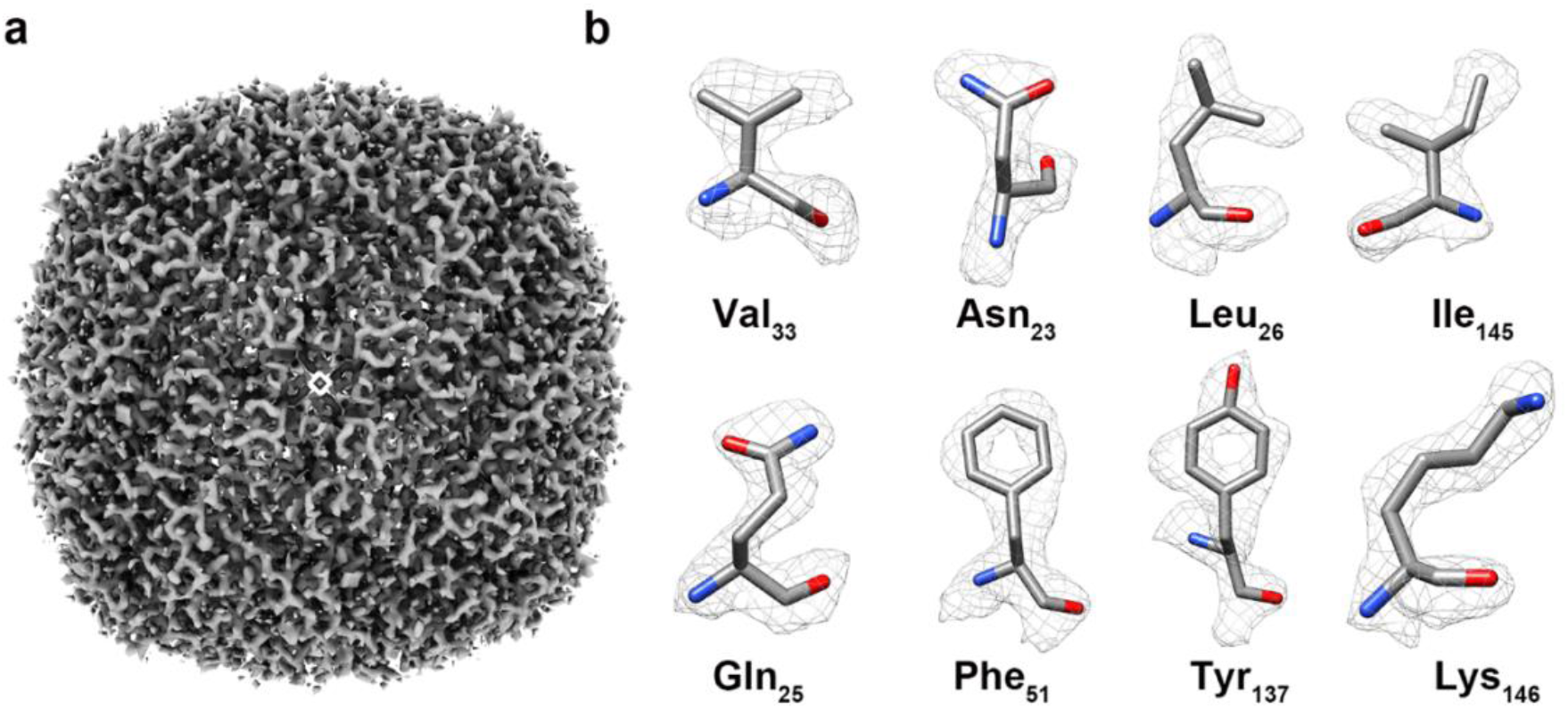
Graphene grids for standard apo-ferritin. **a**, Single-particle reconstruction of apo-ferritin at 2.2 Å resolution. **d,** Representative density maps of selected residues (fitted with PDB model 1FHA) demonstrate convincing side chain structures.

### A 2.6 Å resolution reconstruction of 52 kDa streptavidin with graphene grids

The advantages of using graphene cryo-EM grids can be divided into two categories. For small proteins (<200 kDa), graphene supporting film can effectively reduce the ice thickness without introducing background noise to the images, allowing for higher resolution structural determination of the proteins. For membrane proteins embedded in detergent micelles or liposomes, which are usually hard to acquire in large quantities, graphene grids can improve the protein adsorption, thus overcoming the issues of low protein concentration. Therefore, we evaluated the practical behavior of our graphene grids using a 52 kDa small soluble protein and a membrane protein in detergent micelles and liposomes.

We deposited 52 kDa streptavidin (the smallest soluble protein that has been solved hitherto in EMDB) onto freshly UV/ozone treated graphene grids to prepared frozen-hydrated streptavidin samples in *Vitrobot* with general plunge freezing method. The overall grid montage collected in cryo-EM displays that more than 70% of the grid squares contain thin ice layers that are suitable for data collection (Supplementary Fig. 5a and b). Each grid square possesses a uniform thin ice layer with proteins embedded in (Supplementary Fig. 5c and d), where it is not necessary to screen for good areas in high magnification to save time and efforts. After data acquisition, the 52 kDa streptavidin particles could be identified clearly from the motion-corrected micrographs (Fig. 4a). The good image contrast under such a small defocus value of −0.85 μm further confirms that the ice is much thinner than that in cryo-samples using conventional grids. Furthermore, the first-order reciprocal lattice of graphene could be identified precisely in the Fourier transform of the micrograph (Fig. 4b). The sharp peaks (circled in orange in Fig. 4b) indicate that the raw information contained in the micrograph could reach to at least 2.14 Å in frequency. After single particle analysis using *Relion*^24^ (details in Supplementary Fig. 6), we obtained the structure of apo-state streptavidin at 2.6 Å resolution (Fig. 4c), where the density map has demonstrated convincing structural information of the residues in the beta strands (Fig. 4d). The structure resolution was estimated from gold standard FSC criteria (Fig. 4e) and the quality of the reconstruction was further validated by local resolution and directional 3D-FSC analysis (Supplementary Fig. 7a-d). In addition, the final reconstruction of 2.6 Å structure was obtained from only ~11,000 streptavidin particles with an estimated Rosenthal-Henderson B-factor of 72.8 Å_2_ (Supplementary Fig. 7e), which is much smaller than previously published reconstruction results of streptavidin^25^, indicating that the high-resolution information is better preserved using our graphene grids.

**Figure 4.**
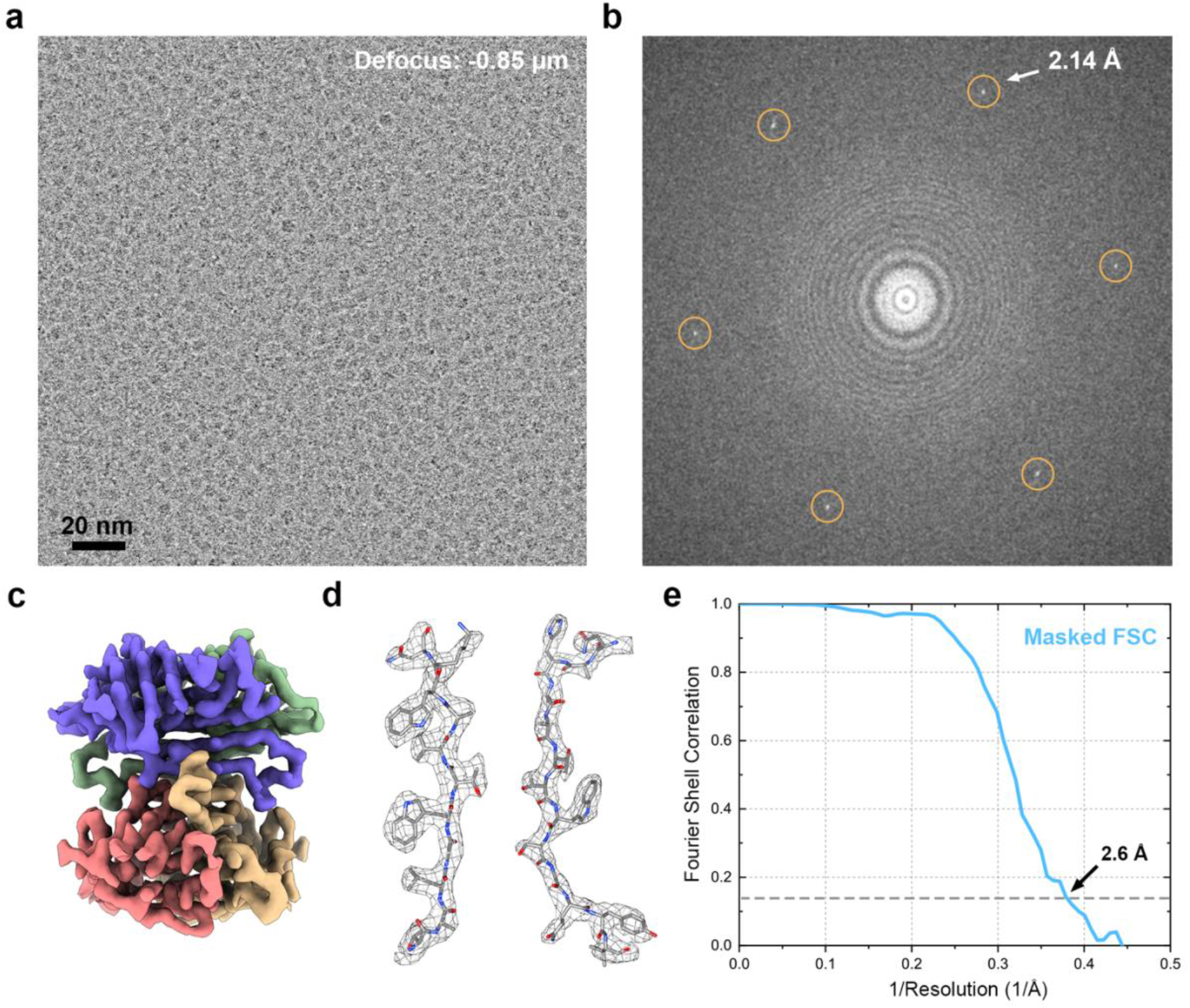
Graphene grids for small soluble proteins. **a,** Cryo-EM micrograph of 52 kD streptavidin particles. Despite a small defocus value (−0.85 μm), streptavidin proteins present nice contrast in the micrograph. **b**, Fourier transform of (**a**) with graphene reciprocal lattice circled in orange. **c,** Single-particle reconstruction of 52 kD streptavidin at 2.6 Å resolution. **d,** Representative density maps of two beta strands (fitted with PDB model 6J6K). **e,** Gold standard FSC (criteria 0.143) curve of the masked map with a reported resolution of 2.6 Å.

### High density of membrane and lipo-proteins on graphene grids

Unlike most soluble proteins, membrane proteins are usually more difficult to obtain in large quantity or high protein concentration. Testing with a bacteria sodium channel (NaChBac) purified in detergent micelles, we observed a five-times improvement of the protein density using our graphene grids (Fig. 5a vs. b). Despite the adsorption of the membrane proteins on graphene supporting film, we observed a reasonable distribution of different views, which is essential for further single-particle data processing. In addition, we reconstitute NaChBac into liposomes for structural investigation, which can provide more orientations of the protein for single particle analysis, as well as reproducing more native physiological environments of the proteins. As the long-time historical challenge to investigate the structure of lipoproteins is the low density of liposomes on a cryo-EM holey carbon grids ^26,27^ (Fig. 5c), our graphene grids remarkably improve the liposome density by providing a hydrophilic surface to attract liposomes (Fig. 5d). These results imply that our graphene grids can assist single particle cryo-EM on membrane proteins in detergent micelles (or lipid nanodiscs) and liposomes by improving the protein density. Our grids will reduce the requirements of cryo-EM sample preparation, allowing for structural investigation of membrane proteins that cannot be produced in large quantity or concentrated to high concentration.

**Figure 5.**
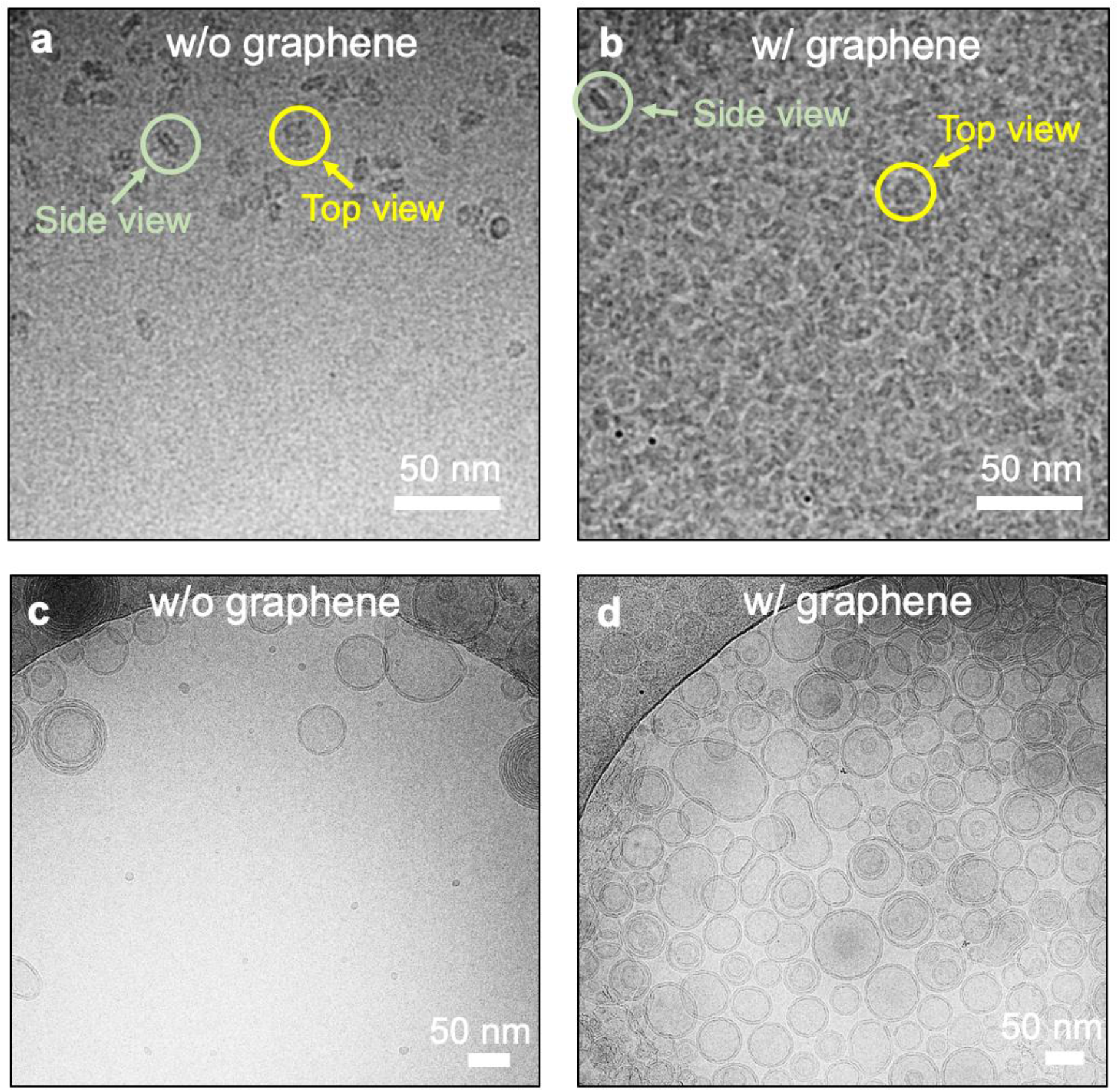
Graphene grids for membrane protein and liposomes. **a,** Cryo-EM micrograph of a bacterial sodium channel (NaChBac) on holey carbon grids. **b,** Cryo-EM micrograph of NaChBac on graphene grids, which increase the protein density by five times. (The protein concentration in solution is 6 mg/mL.) **c,** Cryo-EM micrograph of liposomes on holey carbon grids, where the liposomes prefer to adsorb to the amorphous carbon supporting film. **d,** Cryo-EM micrographs of liposomes on graphene grids. The liposomes adsorbed to the graphene surface uniformly, allowing for cryo-EM data acquisition in the thin ice on graphene over the hole areas.

### Conclusions and discussions

In summary, we developed a robust approach to produce high-quality graphene grids for cryo-EM with about 99% suspended graphene coverage. Our graphene grids provide thinner ice layers and an increased protein density. We have achieved 2.6 Å resolution structure of a 52 kDa streptavidin and a 2.2 Å resolution structure of standard apo-ferritin. For membrane proteins and lipoproteins that are usually hard to prepare in large quantities, we have achieved more than five-fold improvement of protein density, which will aid the studies of membrane proteins in their more native environments. In addition, the method we described to make graphene grids can be easily accessed by most structural biology research groups with reasonable expenses. Our method can also be applied to other nanomaterials such as one-dimensional nanowires and nanotubes, which can allow for more sophisticated grid design targeting specific proteins. We expect our method to benefit the cryo-EM community by improving the sample preparation process.

## Acknowledgement

The work is supported by the Dean for Research Innovation Fund for New Ideas in the Natural Sciences from Princeton University. N.Y. is supported by the Shirley M. Tilghman endowed professorship from Princeton University. The authors acknowledge the use of Princeton’s Imaging and Analysis Center, which is partially supported by the Princeton Center for Complex Materials, a National Science Foundation (NSF)-MRSEC program (DMR-1420541). We thank Mengnan Zhao, Xuelan Wu, Hannah A. Ledford, Ang-Yu Lu, Kibum Kang, Yunfan Guo, Hongwu Qian, Shuai Gao, Xin Zhao, Miaohui Hu, Shiming Lei, etc. for helpful discussion.

## Materials and Methods

### Continuous monolayer graphene synthesis

In this research, commercial copper foils with 99.8% purity (Alfa Aesar #13382) were used as the substrate to grow single-layer graphene by Chemical Vapor Deposition (CVD) method. Before growth, we used nickel etchant (Transene, TFB) to clean the surface of the copper foil. The foil was then placed in a CVD system with a base pressure of 35 mTorr. The foil was annealed in the CVD chamber for 30 min at 1030 °C with a 10 sccm H_2_ flow. Subsequently, single layer graphene was grown at the same temperature (1030 °C) with 60 sccm H_2_ and 3.5 sccm CH_4_ for 30 min to form continuous monolayer. The graphene growth protocol is similar to the one used in ref. ^28^. Alternatively, continuous monolayer graphene on copper foil can be purchased from *graphene supermarket*.

### Graphene grid fabrication

The graphene grown by CVD or alternatively purchased from *Graphene Supermarket* came with graphene on a copper foil. We first coated the graphene on copper foil with MMA EL 6 (or PMMA 495 A2) using a home-made spin coater (See supplementary video). Then the sample was placed in a glow-discharge system backside up to remove the graphene grown on the back side of the copper foil (we used a typical glow-discharge condition: 30s O_2_ or Ar plasma). Afterwards, we used 1M ammonium persulfate (AM) to etch the copper substrate. After the copper foils were entirely gone, we transferred the film (graphene with MMA) to deionized (DI) water 2 times and staying 10 min each time (or you can transfer to DI water once and wait for more than 20 min). After that, we used the Quantifoil (Au 1.2/1.3 300 mesh) to scoop the MMA/graphene film and air dry. Then we placed the grid on a hot plate and bake at 130 °C for ~20 min. Afterwards, we picked up the grid and waited for it to cool down, followed by soaking the grids into acetone vertically. We placed the grids in warm acetone for 30 min to dissolve MMA. Then we transferred the grids to another acetone to further clean MMA. Afterwards, we transferred all grids to IPA to clean off the acetone residue. The process to transfer grids from one organic solvent to another should be very fast to avoid the acetone drying in any case. We took the grid out of IPA vertically and used a filter paper to draw remaining IPA away from the grids. Then we left the grids on a filter paper to air dry them. Afterwards, we placed the grids on a hot plate for another ~20 min to bake off organic residues. The process described above can result in a high yield of suspended, clean graphene films on Quantifoil holey carbon grids. More details can be found in the Supplementary Video. The method was adapted from previous works^29,30^. The materials used in the transfer process are listed below:

- Graphene on Cu was purchased from *graphene supermarket*.
- MMA EL 6 (Item# M310006 0500L1GL) and PMMA 495 A4 (Item# M130004 0500L1GL) was purchased from *MicroChem* (now *Kayaku*).
- Spin coater was made from a computer fan (See Supplementary Video).
- Ammonium persulfate (APS) was purchased from *Sigma Aldrich*.
- Quantifoil TEM grids were purchased from *EMS*.

### Ultraviolet/ozone

We used an ultraviolet/ozone cleaner (UVOCS T10X10 system) to treat graphene surfaces and make graphene grids hydrophilic. This tool is commonly used for cleaning of wafers contaminated with organic substrates. The UV/ozone used a low pressure, quartz, mercury vapor lamp to generate 185 nm and 254 nm UV light. The process took place at room temperature in 10 minutes. A temperature rise was observed upon prolonged cleaning. The surface-treated graphene grids were used for cryo-sample preparation on the same day. An exposure of the surface-treated graphene grids to the air for more than a day may introduce surface contamination or broken graphene that reduces the yield.

### Streptavidin cryo-EM sample preparation

Streptavidin stock solution (1 mg/ml, New England Biology, N7021s) was diluted to 0.5 mg/ml using DI water. UV/ozone freshly treaded graphene grids (Quantifoil 300 mesh Au R1.2/1.3) were used to prepare cryo-samples. In Vitrobot Mark IV (Thermo Fisher), 4 μl 0.5 mg/ml sample was applied to grid, then waited 30s before the blotting. The blot time was 4s with a blot force of 0 in the Vitrobot. After blotting, the grid was rapidly plunged into liquid ethane for the vitrification.

### Data collection

A total number of 1,086 raw movie stacks were automatically collected in 12 hours by SerialEM 3.7 on a 300 kV Cs-corrected Titan Krios using a K2 Summit detector (with GIF Bio-Quantum Energy Filters, Gatan). We collected the raw movies in K2 counted mode at a magnification of 215,000 (in EFTEM mode, spot size 6, C2 aperture 70 μm) with a pixel size of 0.536 Å. The total exposure time was set to 2.4 s with a 0.075 s frame time to generate 32-frame gain normalized mrc stacks. The total dose for a stack is 49 e-/Å2.

### Data processing

Movies stacks were motion corrected using Relion’s own interpretation with a 5 x 5 patch and a 2-fold binning. Full frame CTF values were estimated from non-dose weighted images by CTFFind4.1^31^ in Relion with exhaustive searching. Particles were automatically picked with the Laplacian-of-Gaussian method in Relion Auto-picking. After the particle extraction, the particle stacks were applied a 120 Å high pass filtered (using relion_image_handler --highpass 120) for the following processes. Multiple rounds of 2D classification, 3D classification, 3D auto-refine, CTF Refinement and Bayesian polishing were performed in Relion 3.0 with standard procedure. Density maps are prepared by UCSF Chimera^32^ and Chimera X^33^.

### Data availability

Data which supporting the findings of this manuscript are available from the corresponding authors upon reasonable request. The accession number for the EM map of streptavidin reported in this paper is EMD-20907. The raw data was deposited at EMPIAR-10335.

---

**Supplementary Figure 1.**
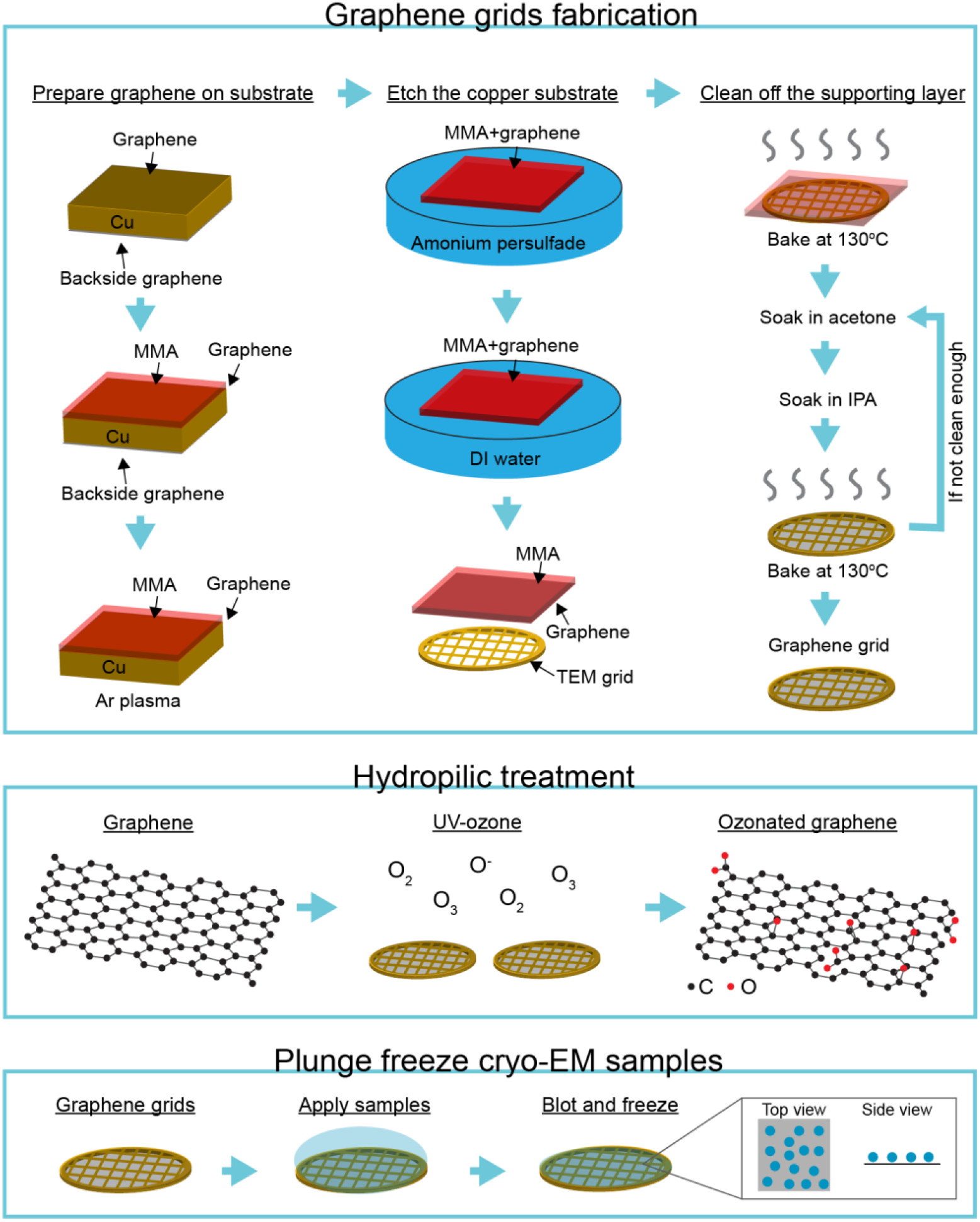
Overview of the method. **Top panel,** Schematic of the process to make monolayer graphene grids for cryo-EM. The continuous monolayer graphene was synthesized on a copper substrate via chemical vapor deposition (CVD). Alternatively, the CVD graphene can be purchased from *Graphene Supermarket*. **Middle panel,** Schematic of surface treatment by UV/ozone. The ozonation of graphene adds oxygenated functional groups to the graphene surface, which makes graphene hydrophilic. **Bottom panel,** Schematic of cryo-EM sample preparation on graphene grids. The plunge freeze process was done by widely used *Vitrobot* by *ThermoFisher Scientific*.

**Supplementary Figure 2.**
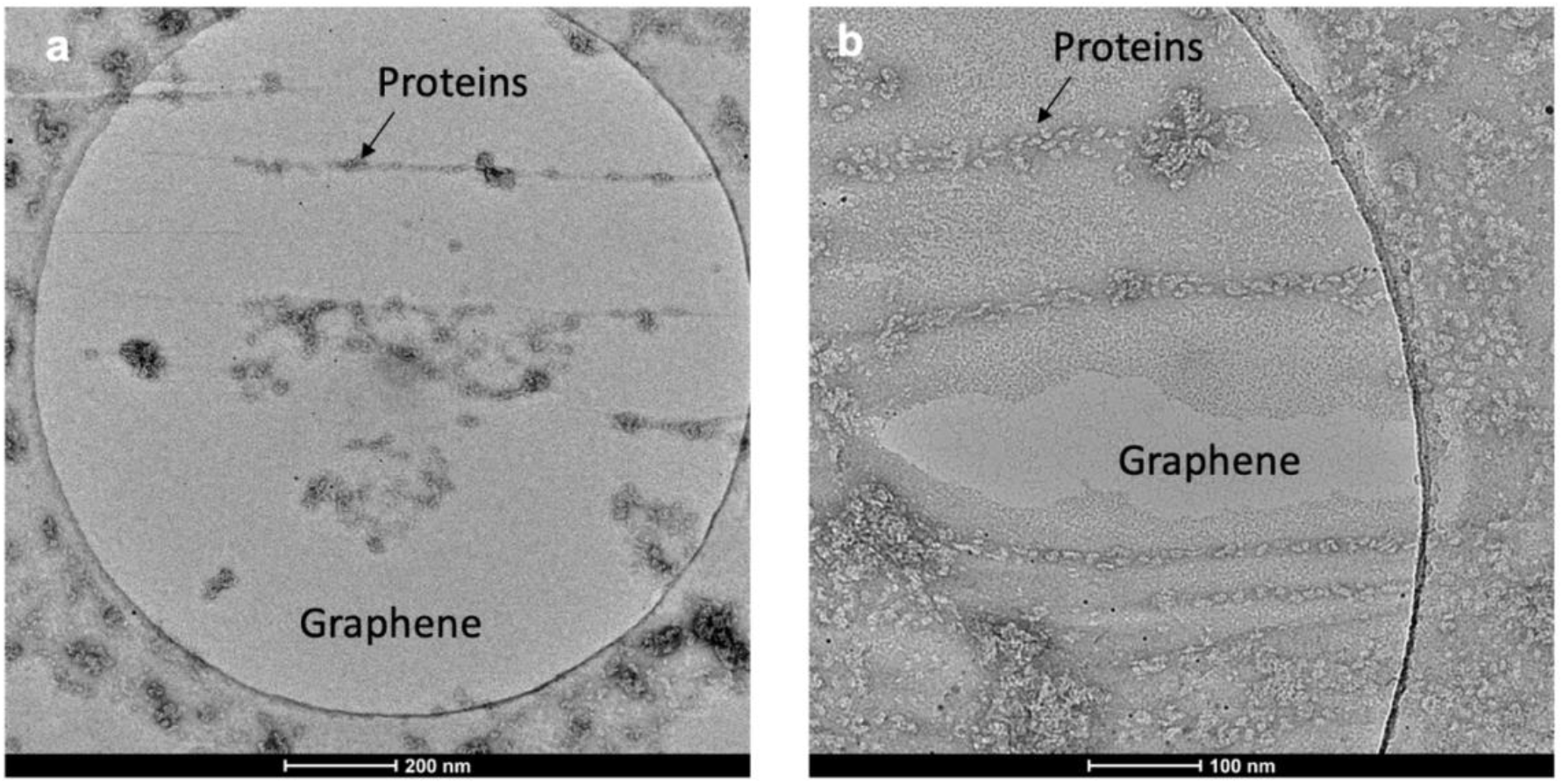
Hydrophobic graphene causing no adsorption or protein aggregations. **a,** TEM image of negatively stained proteins (NaChBac) on a suspended graphene film. **b,** A zoomed in TEM image of negatively stained proteins on graphene. Since graphene is hydrophobic, only a few areas adsorb proteins (indicated in the figures).

**Supplementary Figure 3.**
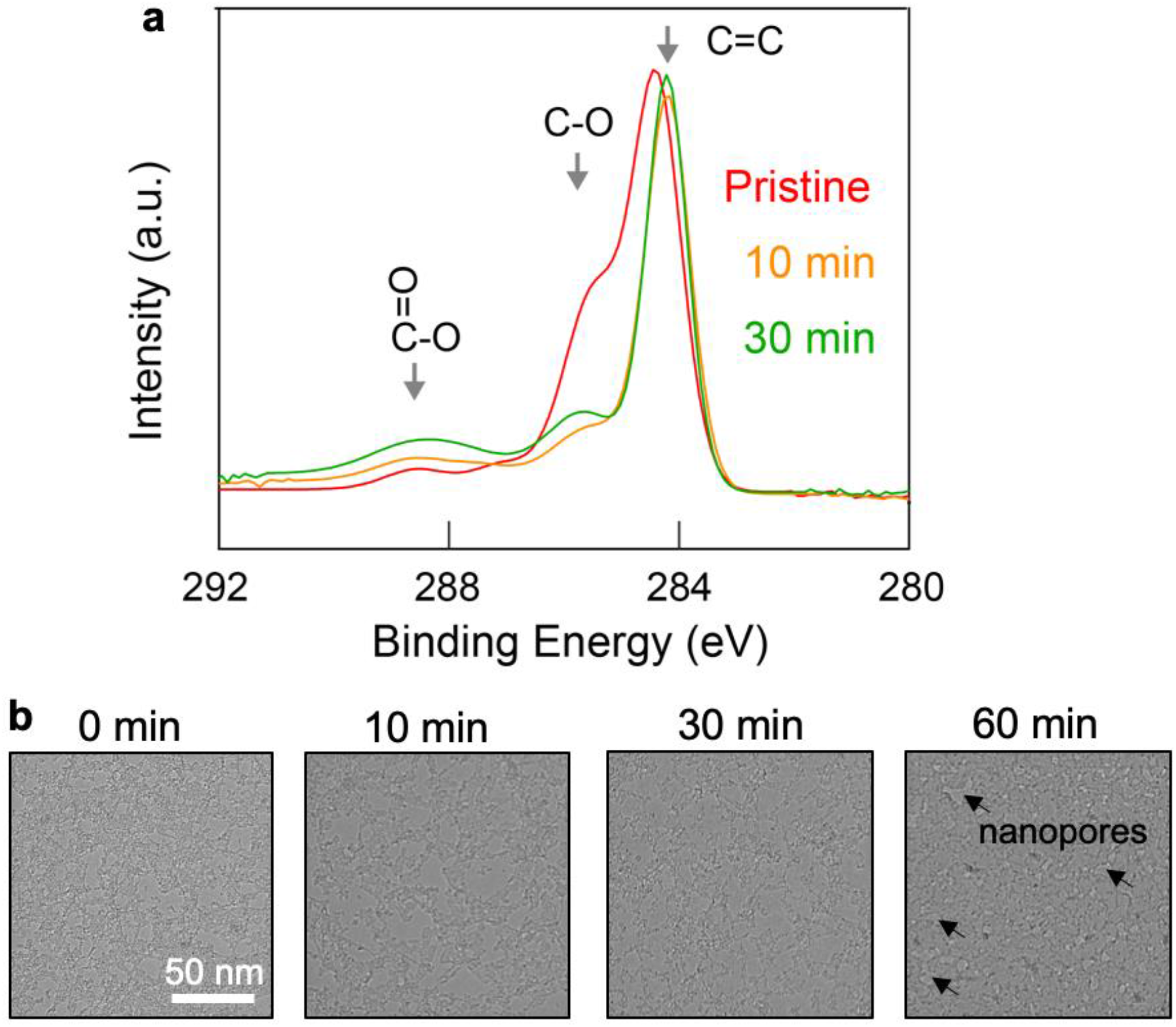
XPS characterization of ozonated graphene surface. **a,** The XPS plots show that in the first 10 min, ozonation helps to clean off the surface contaminations on graphene (indicated by the drop in C-O groups). Further treatment of UV/ozone introduces more C-O and carboxylic groups (-COO) on the graphene surface to make it hydrophilic. **b,** TEM images show that a 10 min UV/ozone can remove some surface contaminations. A 30-min UV treatment further reduced the surface contamination. However, we observed more broken graphene films in the holey carbon areas. A 60-min UV treatment broke most suspended graphene, while the remaining suspended ones produced nanopores.

**Supplementary Figure 4.**
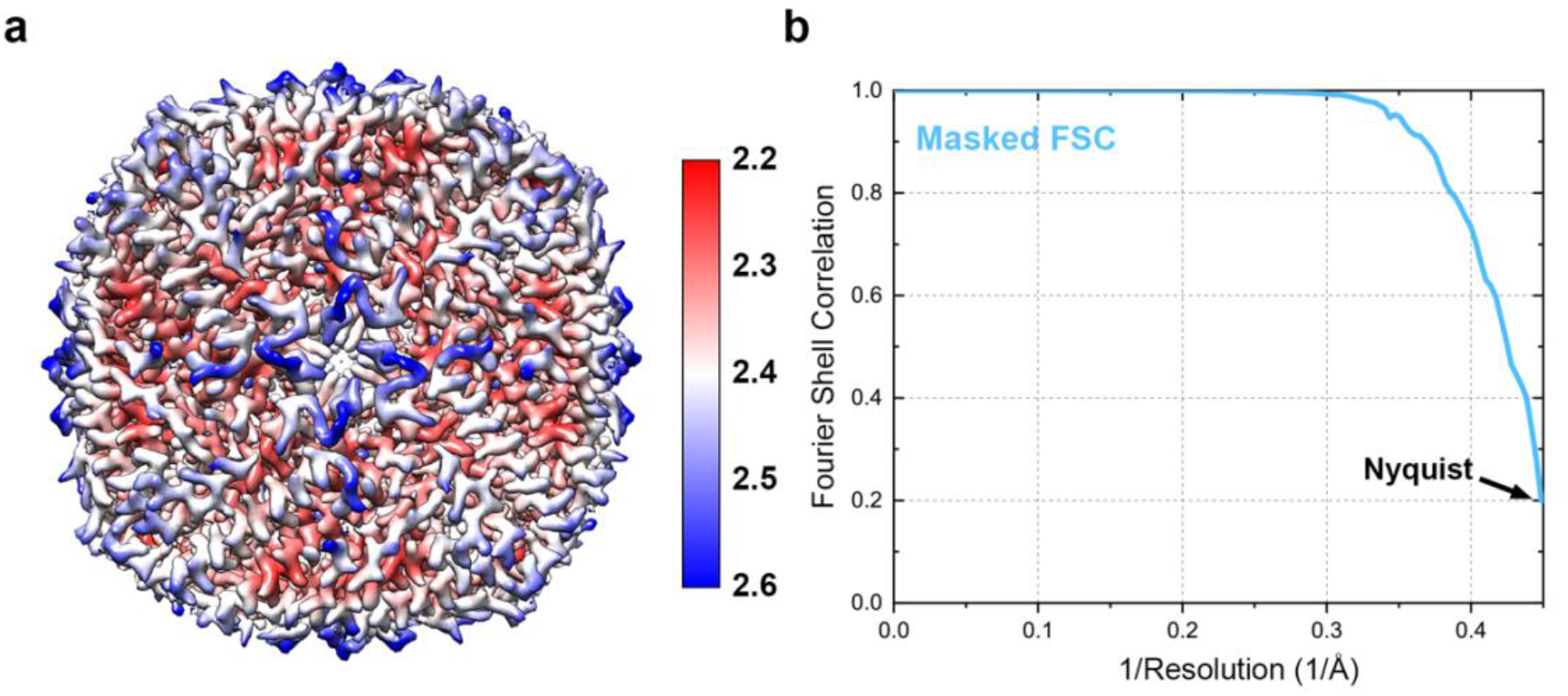
Single-particle reconstruction of apo-ferritin at 2.2 Å resolution. **a,** Resolution map of the apo-ferritin structure in Fig. 3a. **b,** Fourier shell correlation (FSC) plot implying a ~2.2 Å resolution of our structure using the gold standard FSC (0.143).

**Supplementary Figure 5.**
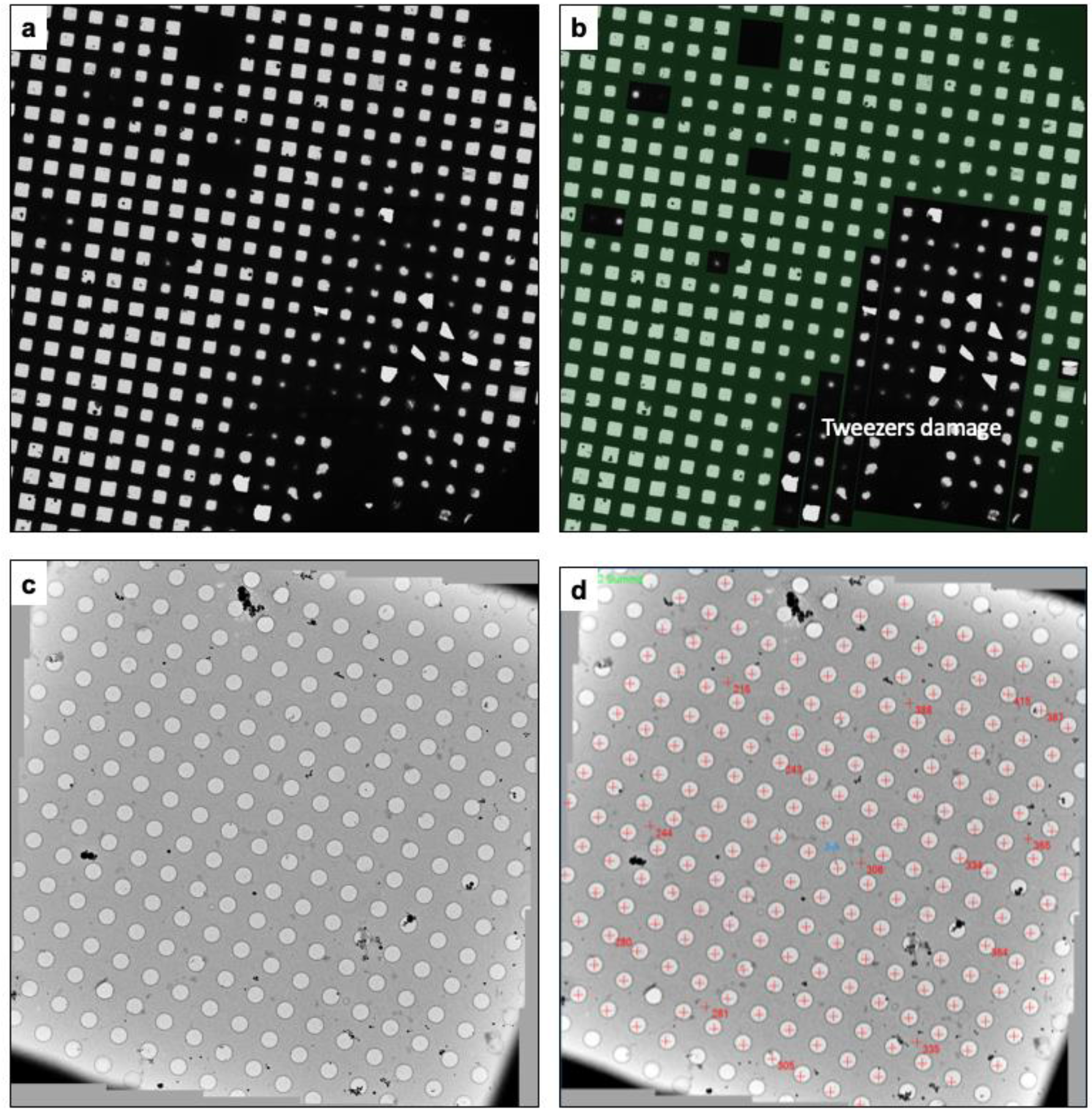
Large-scale montages of cryo-sample using graphene grids. **a,** Grid montage of the cryo-sample prepared in Vitrobot using a graphene grid. **b,** The same grid montage image with effective areas for data acquisition highlighted in green. **c,** Square montage with uniform and thin vitrified ice containing protein particles. **d,** The same square with holes for data collection labeled in red.

**Supplementary Figure 6.**
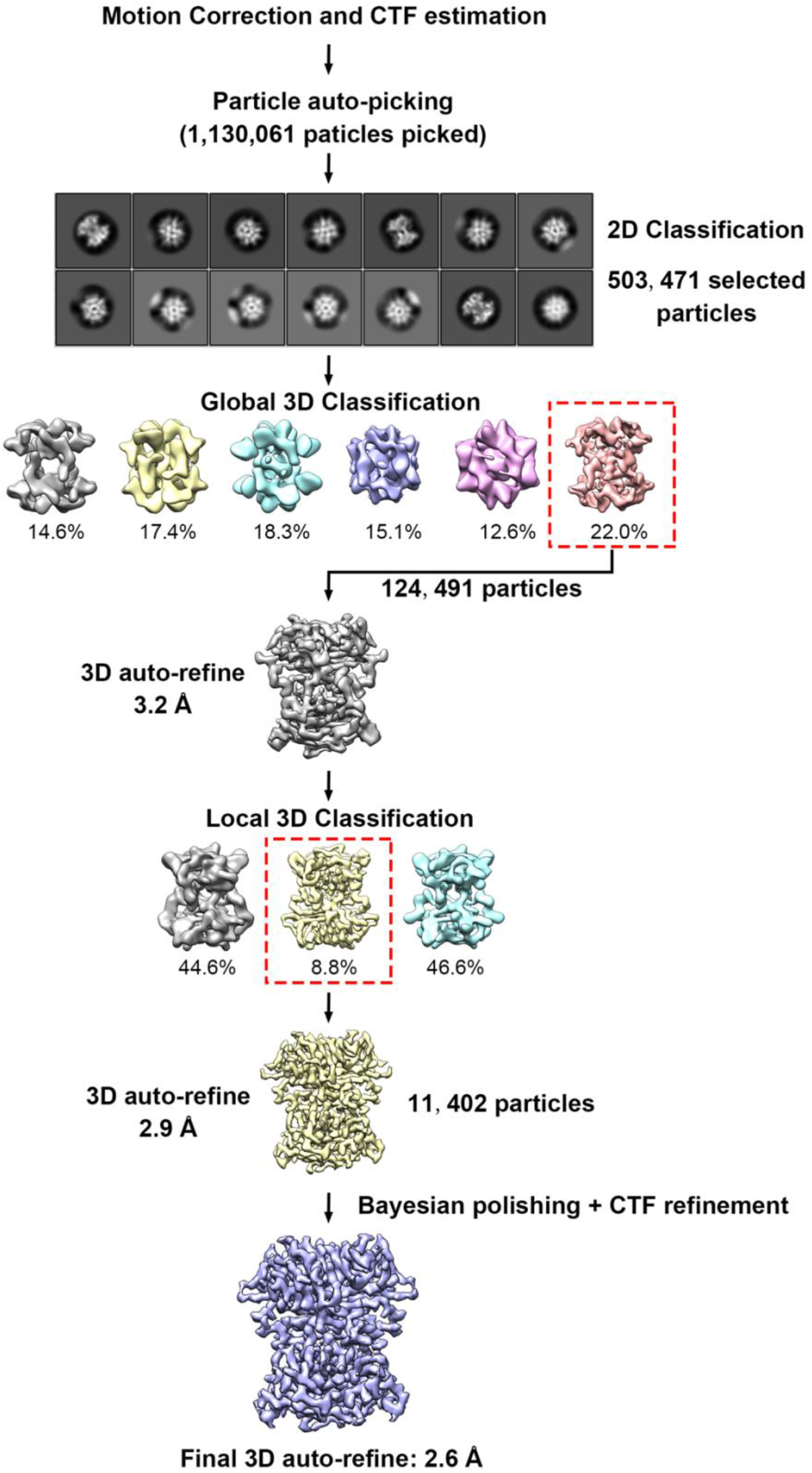
Data processing flow.

**Supplementary Figure 7.**
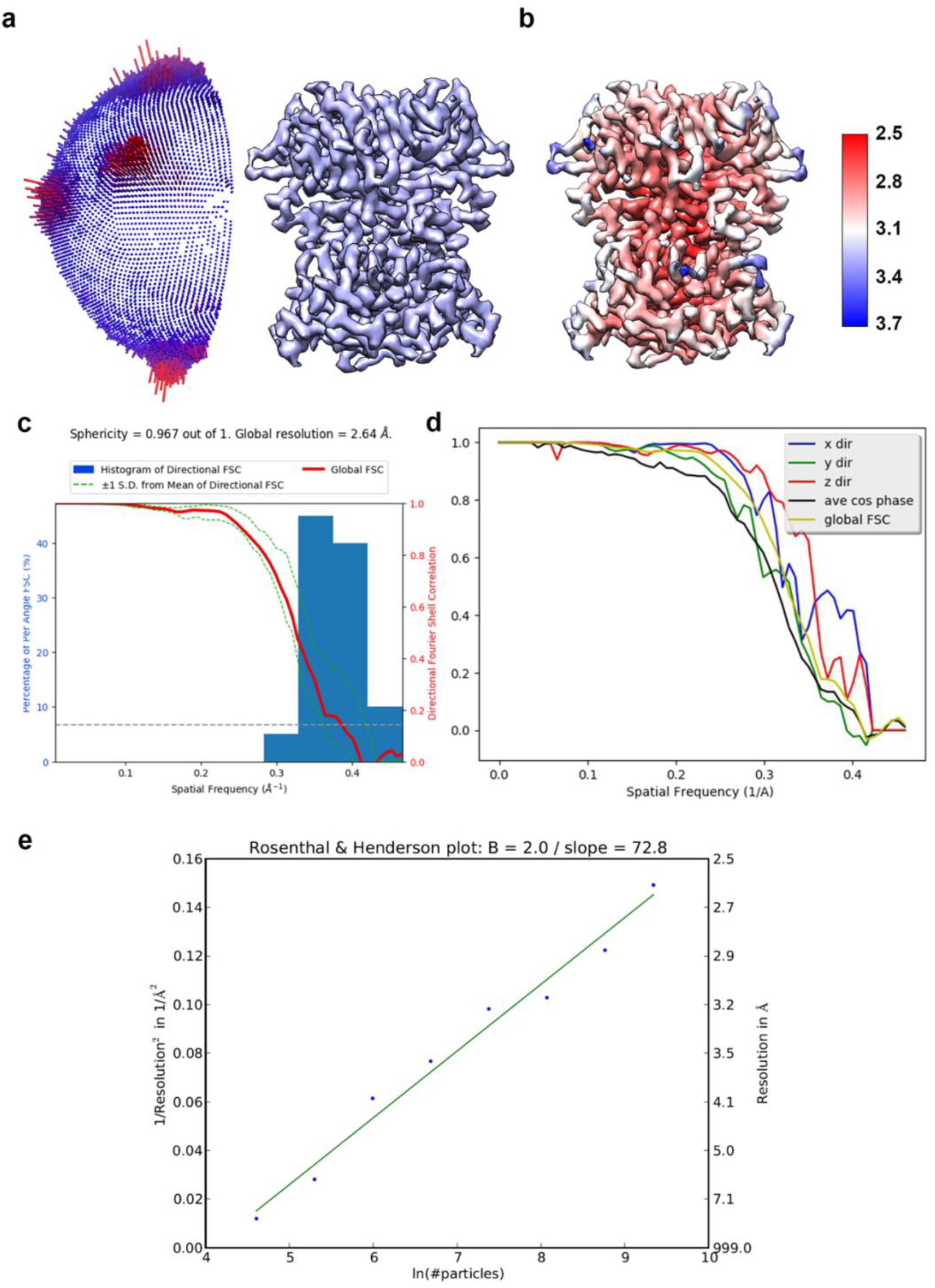
The validation of streptavidin reconstruction. **a,** The angular distribution of streptavidin reconstruction. **b,** The local resolution map. **c,** The histogram of directional 3D-FSC. **d,** Directional 3D-FSC curves. **e,** Rosenthal-Henderson B-factor plot of the streptavidin dataset, with estimated B-factor of 72.8 Å_2_.

